# Design of DNA Aptamers for Lyme disease Diagnosis Combining experimental and numerical approaches

**DOI:** 10.64898/2026.05.13.724892

**Authors:** El Mehdi Issouani, Hugo Da Ponte, Mickael Guerin, Séverine Padiolleau-Lefèvre, Irene Maffucci, Miraine Dávila Felipe, Ghislaine Gayraud

## Abstract

Aptamers are single stranded DNA or RNA molecules selected for their high affinity and specificity to bind target molecules, similar to antibodies. They are commonly selected through the SELEX process, which involves the iterative exposure of a random sequence library to a target and retaining the sequences showing good binding properties. To improve Lyme disease detection, we propose designing aptamers that specifically bind to the *CspZ* protein on the surface of *Borrelia burgdorferi*, the bacterium responsible for the disease.

Starting with a SELEX process consisting of thirteen rounds, from which selected *in vitro* sequence candidates have emerged, we aim to propose a holistic process that selects *in silico* new sequence candidates that are further validated experimentally. Our approach relies on 1) using Machine Learning (ML) techniques, specifically a Restricted Boltzmann Machine (RBM), to digitally replicate the last round of the SELEX process, 2) integrating insights from text analysis methods, such as word2vec and n-grams, into the RBM model trained on the final-round SELEX dataset to represent and compare newly generated sequences with *in vitro* candidates, 3) selecting *in silico* sequences with strong potential to bind to *CspZ* protein, 4) experimentally validating the selected *in silico* sequences of step 3.

Our holistic approach combines biological insights with statistical models to improve the efficiency and outcome of the SELEX process. We enhance the RBM model, designed to replicate the distribution of the final SELEX round, by integrating geometric representations of sequences, which is especially advantageous when dealing with limited datasets relative to the vast sequence space. In addition, it provides *in silico* sequence candidates with strong binding properties.

## 1. Introduction

Aptamers are single-stranded nucleic acid sequences (ssNAs), selected for their high affinity and specificity in binding to target molecules, much like antibodies. They are identified through an experimental protocol called *Systematic Evolution of Ligands by EXponential enrichment* (SELEX), which allows isolating aptamers with desirable binding properties from large random libraries [1, 2]. These versatile ligands are valuable in diagnostics, drug development, and research, binding effectively to proteins, small molecules, and even complex targets such as cells [3]. Aptamers can act as precise tools for targeting biological functions and aid in the recognition and interaction with drugs or chemical compounds [4]. *In vitro* selection has enabled the discovery of specific RNA and DNA molecules, including ribozymes and diagnostic/therapeutic aptamers. SELEX-developed aptamers can control protein activity with light-sensitive groups, regulating biological processes, as demonstrated with thrombin and cytohesin-1 [5] as demonstrated with thrombin and cytohesin-1 [5], enabling the development of unique RNA and DNA aptamers for use in diagnostic kits [6, 7] and therapeutic applications [8].

SELEX involves exposing an initial random sequence library to a target molecule, only retaining sequences that specifically bind to the target, and eliminating non-binding ones. This process is repeated several times, through what is called *rounds*. The remaining sequences at the end are potential candidates for binding to the target (see [9]). Acquiring sequences, aptamers, or target molecules for conducting the SELEX protocol involves considerable costs, while the resulting database remains limited compared to the complete sequence space (10^13^-10^15^ instead of 10^23^ for sequences of length 40). Consequently, researchers are increasingly adopting computational methods to analyze SELEX data and enhance its utilization [10]. More recently, SELEX–ML integrations have been used to explore RNA fitness landscapes and predict sequence evolution [11]. Similarly, integrative computational strategies are increasingly applied in aptamer-based nanomedicine [12].

Current serological methods for Lyme disease diagnosis remain debated, highlighting the need for more direct and specific detection strategies. In this context, an integrative approach is developed, to design DNA aptamers capable of identifying the presence of *Borrelia burgdorferi* (*Bb*) bacteria characterized by its surface protein *CspZ*. To this end, the proposal consists in the combination of experimental and numerical approaches, including *in vitro* experimental data (SELEX) and machine learning technics, to identify aptamers that can bind to the surface protein *CspZ*.

The most common approach developed to identify ssNAs with specific binding properties relies on characterizing their secondary (i.e., their base pairing pattern) and tertiary (i.e., their 3D organization) structures, which play a key role in molecules function. This characterization, known as *folding*, can be obtained either experimentally or by (experimentally-driven) computational prediction. Experimental protocols (such as crystal structure analysis or high-resolution spectroscopy) yield the structural data with the highest accuracy. However, these methods remain time-consuming and costly. On the other hand, *in silico* structure prediction, usually derived from thermodynamic principles [13, 14, 15], and more recently from machine learning methods [16, 17], do not provide a sufficiently accurate alternative. We refer to [18] the readers who are interested in recent ML-based advances in RNA structure and interaction prediction. Moreover, the implementation of numerical or experimental folding prediction during SELEX is impractical because of the relatively short time duration of each round compared to the large number of sequences involved and the high level of structural flexibility of ssNAs, which varies depending on the presence or absence of the target.

Given that structural data are either unavailable or only partially accessible, a promising strategy is to work directly with the SELEX sequences by extracting their statistical features and proposing suitable models to approximate their probability distribution. Comparable ML–statistical techniques have improved peptide design from limited data [19]. Following this direction, Di Gioacchino et al. [20] combined the direct exploitation of the primary structure with a Restricted Boltzmann Machine, i.e., a Machine Learning probabilistic model, to enhance SELEX results. Relying on a very large and highly diverse sequence dataset, their method allows for analyzing specific SELEX rounds, predicting aptamer presence and relative frequency in later rounds, and generating new candidates absent from the training set. Furthermore, their new predicted sequences were experimentally validated and showed strong binding to their target (thrombin). They demonstrated the performance of their approach in the context of the availability of a large dataset that has a high diversity and more reference metadata.

In the context of a more moderate dataset size, Di Gioacchino et al.’s [20] approach can be extended and complemented by deriving a data representation directly from their model. Inspired by Natural Language Processing (NLP) methods like *n*-grams and Word2Vec [21, 22], we explore how SELEX sequences can be embedded as numerical vectors in a latent space. To evaluate the relevance of these embeddings, we rely on experimental data from thirteen SELEX rounds targeting the *CspZ* protein, among which 16 aptamers were selected and experimentally tested individually for their binding affinity. The aim is to generate new sequences that resemble these validated examples, which requires defining suitable similarity metrics.

One of the aims of this work is to digitally replicate the final round of SELEX (*in silico*), complementing the experimental (*in vitro*) process. Building on vector-based embedding techniques, stochastic or deterministic, we leverage experimentally derived SELEX data to compare sequences in a latent space and generate new high-affinity binders absent from the current dataset [9], which have subsequently been validated through additional experimental evaluation.

To achieve this, we approximate the unknown distribution of sequences from the final SELEX round using a model that assigns likelihoods to any possible sequence and is used to generate new sequence candidates, allowing to both describe the final-round sequence space and enrich it with promising binders.

Hence, we aim to provide a holistic approach that selects *in silico* new sequence candidates that are further validated experimentally. In particular, we propose a comprehensive procedure by leveraging RBM models to analyze SELEX data, focusing on sequence representations and their relationships. We pre-process the dataset, use RBMs to extract meaningful features, and generate novel sequences. We use appropriate distance metrics to compare sparse representation (i.e. *k*-mers-derived) and dense representation (i.e. RBM-derived) representations, aiming to evaluate how effectively they capture sequence similarities. A subset of the generated sequences was selected based on these representations and similarity metrics, and subsequently validated *in vitro* for their binding affinity, confirming the practical relevance of the proposed approach. Our findings highlight the benefits of applying ML approaches to biology, offering insights into aptamer selection and potential applications in diagnostics and therapeutics.

## 2. Materials and Methods

### 2.1. Dataset

*In vitro* experiments were conducted to isolate favorable aptamers, specifically those capable of binding to the *CspZ* protein through the SELEX method [9].

The SELEX process involved 12 rounds of selection, with around 10^12^ sequences per round. For each round, a random sample of 10^5^ ssDNA sequences are obtained by Next-Generation Sequencing (NGS). Each sequence is 76 nucleotides long, consisting of a central variable region of *d* = 40 nucleotides (the aptamer) bordered on both sides by fixed 18-nucleotide primer sequences (used to stabilize the sequences for PCR amplification; see [9] for more details). The initial random database is balanced, having an overall proportion of 25% of each nucleotide. In what follows, only the aptamer regions (excluding primers) are considered as the sequence dataset.

Table 1 summarizes the data with the counts (in percentage %) of ssDNAs with and without repetitions, and the count of the five most frequent sequences in each round. Note that the most frequent sequence in round *i* is not the same as the most frequent sequence in round *j* when *i* ≠ *j*. This suggests that the selection of potentially favorable sequences occurred during the SELEX process. As the rounds progressed, some sequences disappeared, even those that were initially highly frequent. In contrast, others increased in frequency and eventually became dominant, despite having low frequencies in the initial rounds^1^. In Figure 1B, the decreasing entropy as the rounds advance highlights a clear selection process.

**Table 1.**
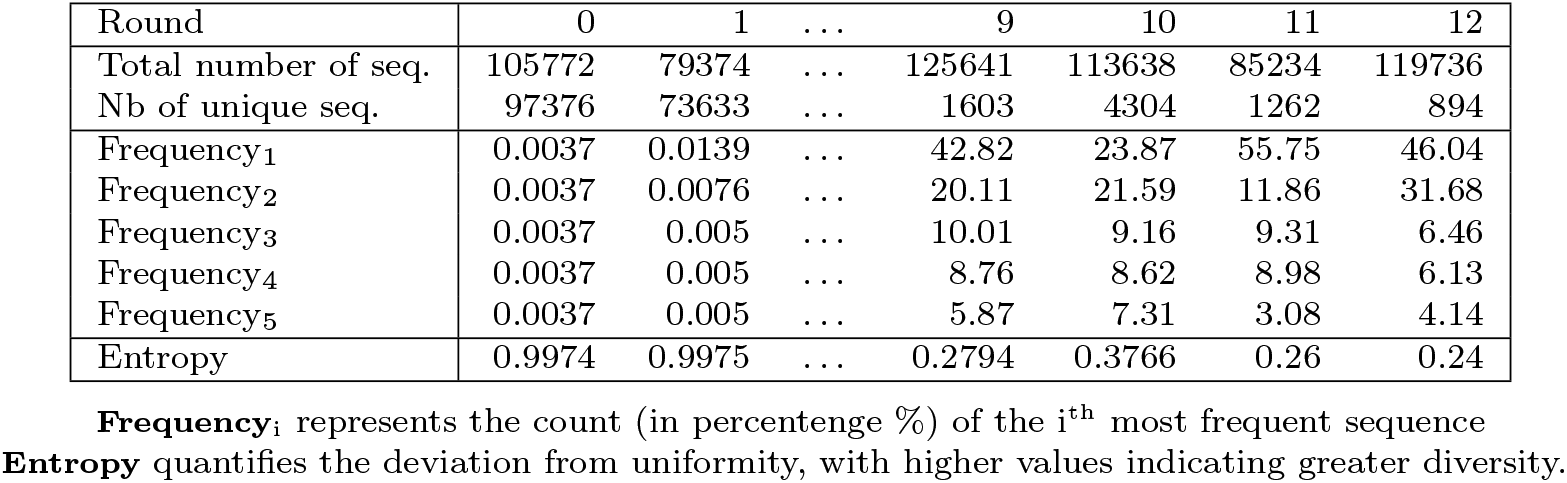
Analysis of NGS sequencing results.

| Round | 0 | 1 | ... | 9 | 10 | 11 | 12 |
| --- | --- | --- | --- | --- | --- | --- | --- |
| Total number of seq. | 105772 | 79374 | ... | 125641 | 113638 | 85234 | 119736 |
| Nb of unique seq. | 97376 | 73633 | ... | 1603 | 4304 | 1262 | 894 |
| Frequency<sub>1</sub> | 0.0037 | 0.0139 | ... | 42.82 | 23.87 | 55.75 | 46.04 |
| Frequency<sub>2</sub> | 0.0037 | 0.0076 | ... | 20.11 | 21.59 | 11.86 | 31.68 |
| Frequency<sub>3</sub> | 0.0037 | 0.005 | ... | 10.01 | 9.16 | 9.31 | 6.46 |
| Frequency<sub>4</sub> | 0.0037 | 0.005 | ... | 8.76 | 8.62 | 8.98 | 6.13 |
| Frequency<sub>5</sub> | 0.0037 | 0.005 | ... | 5.87 | 7.31 | 3.08 | 4.14 |
| Entropy | 0.9974 | 0.9975 | ... | 0.2794 | 0.3766 | 0.26 | 0.24 |

**Frequency<sub>i</sub>** represents the count (in percentge %) of the $i^{\text{th}}$ most frequent sequence **Entropy** quantifies the deviation from uniformity, with higher values indicating greater diversity.

**Figure 1.**
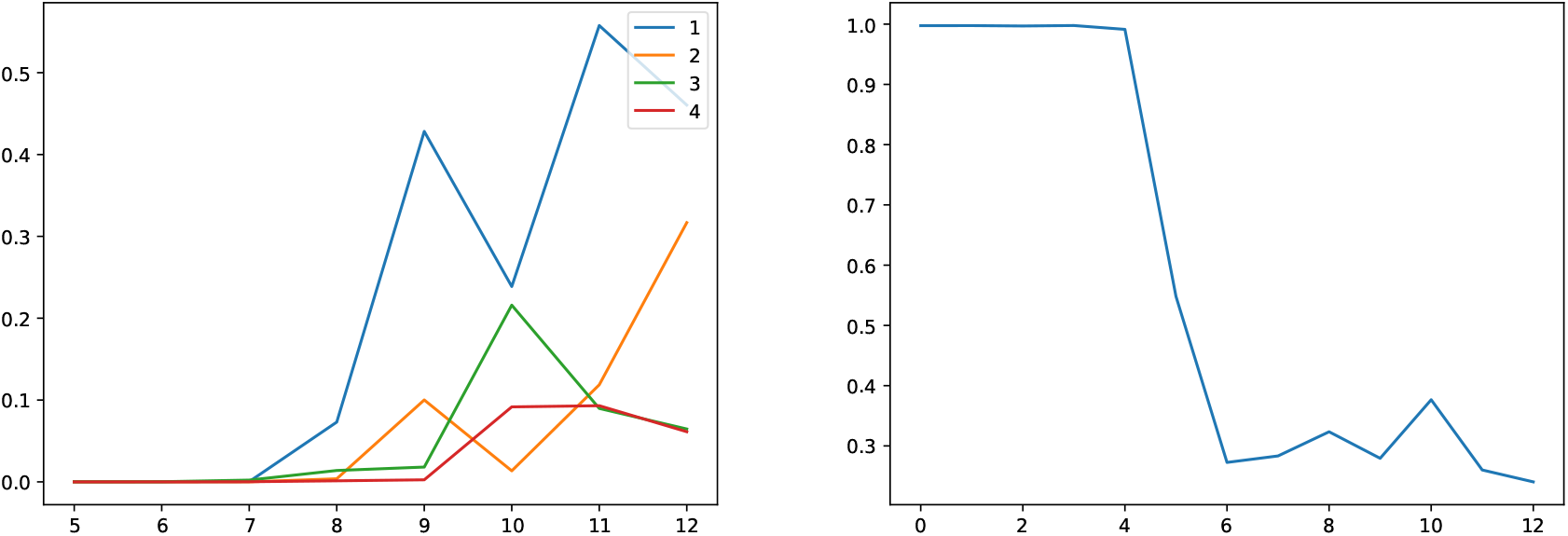
Frequency and entropy evolution versus round number (from round 0 to round 12) Figure 1A. Frequency evolution of the 4 most frequent sequences of the last round Figure 1B. Entropy evolution through different SELEX rounds

Figure 1A illustrates the frequency evolution of certain sequences in the different SELEX rounds, specifically those that are most frequent in the final round. As observed, these sequences started with very low frequencies, which gradually increased over successive rounds until they dominated the final selection, particularly the sequence represented by the blue curve (curve number 1).

The first point to note is that, during the SELEX rounds, the experimental conditions were deliberately varied to reduce selection biases related to the equipment and techniques used. We recall that the target in this study is the *CspZ* protein, a surface protein of the pathogen involved in Lyme disease. *In vivo*, however, *CspZ* is not isolated but embedded in the bacterial membrane of *Bb*, exposing only part of its surface. Early SELEX rounds, performed using the isolated protein, may therefore enrich sequences binding to both accessible and less accessible epitopes. However, only those recognizing epitopes exposed on the bacterial surface are likely to be functionally relevant.

As described in Guérin et al. [9], the selection target was switched to whole bacteria at round 10, thereby reducing off-target binding. This shift is consistent with the slight increase in sequence diversity observed at round 10 in Table 1 and Figure 1B, interrupting the otherwise decreasing trend across rounds.

In this study, we use sequences from round 12 (aptamer regions only) as the dataset to train an RBM model and construct vector representations. Other rounds are used indirectly for three purposes: (1) hyperparameter calibration, (2) model selection, (3) *In vitro* candidates selection.

Among the 16 experimentally tested candidates, 3 are selected based on their high binding affinity to the target. These 3 experimentally tested, validated and selected sequences are referred to as *in vitro* candidate sequences.

The 16 potential candidates are identified from all SELEX rounds based on their frequency, retaining those that appear in at least one round with a frequency above 1.5%. From this subset, only three are selected using additional criteria such as KD interaction strength, cytometry, and the presence or absence of G-Quadruplexes^2^ (G4).

More precisely, flow cytometry allows for the measurement of fluorescence signals associated with the binding of an aptamer (typically labeled with a fluorophore) to its target, on a cell-by-cell or particle-by-particle basis. The aptamer-tagged fluorophore emits a detectable fluorescence signal upon binding. Each particle or cell is individually analyzed, providing precise data on fluorescence intensity for each event. By comparing fluorescence intensities between populations with and without binding, or at different aptamer concentrations, it is possible to estimate the strength of interaction and affinity. This approach enables the generation of binding curves and the determination of key parameters, such as the affinity constant (Kd). (For the full process, see Supplementary Information (SI) Appendix, section 1).

By intersecting the sequences that passed all experimental validations, the retained *in vitro* candidate sequences are called **A6, A9, and A10**, with A9 and A10 showing the best results. These three sequences will guide our investigation into new sequences that are potential candidates, say *in silico* candidates.

In round 12, a small number of sequences dominate, appearing in more than 50% of cases, significantly skewing the results. Therefore, taking into account the specialization of the SELEX sequences as the rounds progress and due to the model validation (see Section 2.2), we use unique sequences from round 12, disregarding their repetitions.

**To summarize:** In this study, our dataset is derived from Round 12 of the SELEX experiment, which contains 894 unique sequences. After removing invalid sequences (e.g., sequences containing ‘N’ for non-identified or missing characters, or sequences with a length different from 40), the number of unique sequences without repetition is reduced to 642. For training and evaluation, the dataset is randomly divided into a training set and a test set, with 578 sequences (90%) assigned to the training set and 64 sequences assigned to the test set.

### 2.2. RBM model

To replicate numerically the last round of SELEX, we use the RBM as a probabilistic model on the sequences, since it allows for capturing their internal dependency by the introduction of hidden variables. Once an RBM is fitted on the sequences ***v*** = (*v*_1_, …, *v*_*d*_) of our dataset with ***v*** ∈ 𝒮 = {*A, C, G, T*}^*d*^, it is used to generate new sequences.

Let ***V*** be a random vector of size *d* whose realization is ***v***, with an unknown probability distribution, that we model by an RBM which consisting on a parametric family of joint probability models {ℙ_*θ*_; *θ* ∈ Θ} over the visible variables ***V*** and latent variables ***H*** of size *d*_*H*_ and realization *h*, each defined via a Gibbs distribution as follows

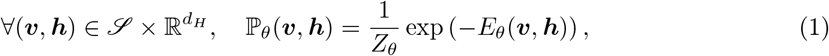

where *E*_*θ*_(***v, h***) represents the energy function, *Z*_*θ*_ the partition function that normalizes the distribution, and *θ* the set of parameters. The marginal distribution over ***V***, obtained from ℙ_*θ*_, is then used to approximate the distribution of ***V*** (if ***H*** is continuous, the sum is replaced by the appropriate integral)

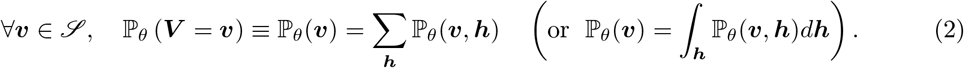

A key feature of an RBM is the factorization of its conditional distributions

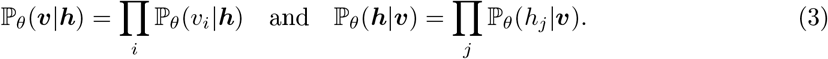

This decomposition not only enhances flexibility but also allows parallel training.

Following the approach proposed by Di Gioacchino et al. [20], we use an energy function that incorporates a double Rectified Linear Unit (double ReLU) represented below by 𝒰_*j*_ (*h*_*j*_). The energy function is defined as

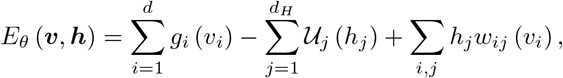

where *θ* = {(*g*_*i*_, 𝒰_*j*_, *w*_*ij*_) |(*i, j*) ∈ ⟦1, *d*⟧ × ⟦1, *d*_*H*_⟧}, and the dimension of the set of parameters is dim(*θ*) = 4 × *d* + 4 × *d*_*H*_ + 4 × *d* × *d*_*H*_ (the first and third multiplicative factor 4 are due to the four nucleotide categories since there is one parameter for each base type, and the middle comes from the double RELU coefficients). A key advantage of this formulation is that the log-likelihood log ℙ_*θ*_(***v***) of a sequence ***v***, obtained by marginalizing the joint distribution ℙ_*θ*_(***v, h***) over the hidden variables ***h***, has an explicit analytical expression. Figure 2 is a graphical illustration of the architecture of such a model.

**Figure 2.**
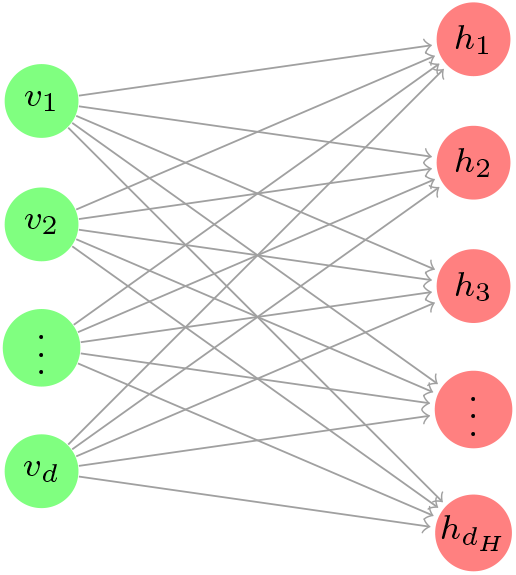
Architecture of an RBM that has d visible nodes and d_H_ hidden nodes

In practice, given the training data ***v***^(1)^, …, ***v***^(*T*)^ of size *T*, the goal of inference is to determine the optimal parameter 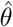 by minimizing the average negative log-likelihood

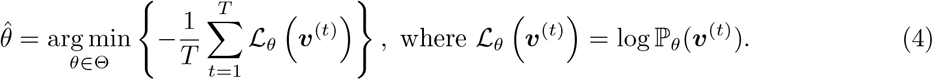

To find the optimal parameter 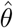, we employ stochastic gradient descent, iterating over the training data by selecting one data sample ***v***^(*t*)^ at each step.

In the RBM case, Gradient Descent consists of iteratively updating the parameter *θ* as follows

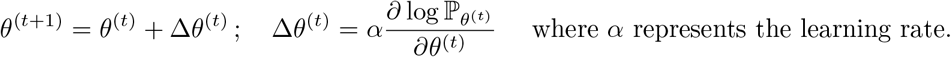

For a model of the form (1) to (2) depending on a parameter *θ*, the gradient of the negative log-likelihood given a single training example ***v***^(*t*)^ is given by

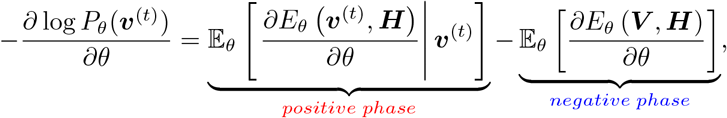

which requires a numerical approximation. Estimating the positive phase term is made by generating ***h*** given ***v***^(*t*)^ using the model. However, estimating the negative phase term is a more difficult task since it requires computing a double sum over ***v*** and ***h***. Instead, this double expectation is approximated using MCMC techniques [23, 24] and Contrastive Divergence as in Hinton (2002) [25]; it consists in replacing the full expectation with a synthetic point estimate at 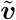 obtained through Gibbs sampling which starts at ***v***^(*t*)^ (see Subsection 2.1 of SI Appendix for a detailed explanation of the Contrastive Divergence algorithm).

Consequently, two types of samples are required:

1. 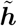 given ***v***^(*t*)^ (∼ *P*_*θ*_(***h***|***v***^(*t*)^)) (*positive phase*) yielded from the right equation in (3).
2. 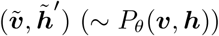 (*negative phase*) yielded from the left and right equations in (3).

The two steps are repeated iteratively until the Markov chain converges to the stationary distribution, which guarantees an unbiased estimate of the gradient.

#### Model Selection and validation

The main trade-offs and hyperparameters considered are: (1) The size of the hidden layer (*d*_*H*_), (2) Whether to account for sequence repetitions (by including sequence counts), (3) the number of iterations.

To select the best hyperparameters, various configurations are tested. The dataset is randomly split into 90% for training and 10% for testing as mentioned above.

For the number of hidden nodes *d*_*H*_, a grid of values {30, 40, …, 120} is used. For each value, the average log-likelihood across test sequences is calculated. This is combined with the number of iterations (point (3)): for each hidden layer size, multiple models are trained with varying numbers of iterations. The optimal number of iterations is chosen based on the highest average log-likelihood for the corresponding hidden layer size. Similarly, two models are compared: one that accounts for sequence counts and one that does not. The one without repetition outperforms the one with repetition (in terms of the matching between training and test datasets and their comparisons with uniformly distributed data in 𝒮).

We fix the number of updates and the batch^3^ size, then the number of epochs^4^ (or the number of iterations) is obtained by dividing the number of updates by the number of batches in the training dataset *N*_*B*_, where *N*_*B*_ = *Training dataset size/Batch size*.

### 2.3. Representations and distances

Before identifying *in silico* candidate sequences, it is necessary to define an appropriate vector representation and to select suitable distance measures that will guide their comparison and selection, in order to detect *in silico* candidates that are close to the *in vitro* candidates.

#### Representations

One of the key steps in applying statistical methods to unstructured data is to convert or encode it into a numerical format, such as a number, vector, matrix, or tensor. This numerical transformation, which allows for the description or characterization of the object, is known as a representation. In ML, such representations are often referred to as embeddings. There are two primary strategies for obtaining them. A stochastic approach involves training a neural network on a task, where intermediate layers learn optimized, low-dimensional representations, called dense embeddings, as seen in methods like Word2Vec or Doc2Vec. A deterministic approach directly computes features from data, such as by counting *k* -mers or detecting specific patterns, resulting in high-dimensional, sparse embeddings. We now outline the sequence representations employed in this work:

– *Raw representation* The simplest form of representation is the sequence itself.
– *Dense representation (continuous)* From an RBM model, a vector ***h*** sampled from 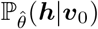 is random and varies between runs. To reduce this variability and obtain a more stable representation of the sequence ***v***_0_, we use the mean vector 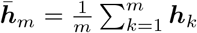, where each ***h***_*k*_ is independently drawn from 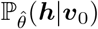. This average vector serves as a dense (continuous) representation of the sequence ***v***_0_.
– *Sparse representation (discrete)* Inspired by NLP, where a document is typically represented as a sparse vector encoding *n*-gram frequencies over a fixed vocabulary, we represent aptamer sequences using sparse *n*-gram vectors. This representation captures sequence patterns and structural motifs, offering practical advantages and interpretability [26, 27]. In aptamer analysis, *n*-grams are referred to as *k*-mers (subsequences of length *k*, commonly used in sequencing) and yield sparse (discrete) representations by counting their occurrences in each sequence.

#### Distances

Having defined both raw and representation-based forms of sequences, we now introduce some distance metrics to quantify their similarity. These distances enable ranking sequences by proximity to a given *in vitro* candidate, supporting structured comparison and analysis. See [28] for a comprehensive overview of distance metrics, including Levenshtein, Hamming, and others in biological contexts (see also [29]). In the following, we present the metrics we consider:

– *Hamming and Levenshtein distances* Both allow for measuring dissimilarity between sequences. Hamming counts the number of substitutions between strings of equal length, while Levenshtein computes the minimum number of edit operations (insertions, deletions, and substitutions) required to transform one string into another, allowing comparison of strings with different lengths [30, 28, 29]. We apply Levenshtein distance to raw sequences.
– *Cosine similarity and ℓ*_2_*-distance* Cosine similarity is widely used in text mining and data analysis due to its efficiency in high-dimensional spaces and its independence from vector magnitudes. This metric is suitable for sparse representations and is defined by,

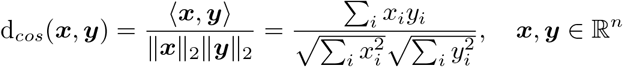

where ⟨***x, y***⟩ is the Euclidean dot product, and ∥***x***∥ _2_ the *Euclidean norm* with *n* being the dimensionality of the space, potentially large.

For dense vectors of small dimensionality, such as those produced by models like Word2Vec or RBM, the *ℓ*_2_ distance is more appropriate, as it captures absolute differences in continuous values and is sensitive to the actual spatial positions of the points. It is defined by

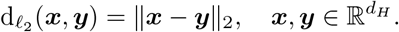

## 3. Results

Once trained, the RBM model is used both to evaluate the log-likelihoods of sequences and to generate new sequences, which are then examined and filtered through the broader analysis described in the following.

### 3.1. Empirical model validation

To validate the model empirically, we first compare the log-likelihood distributions of the training, test of our dataset and random^5^ sequences. The test set closely matches the training distribution, while random sequences show much lower log-likelihoods, indicating that the model captures relevant statistical features of the data and generalizes well.

Figure 3 illustrates these results. The first plot (Fig 3a) shows histograms of log-likelihoods for the training (blue), test (orange), and random (green) sequences. The second plot (Fig 3b) highlights the log-likelihoods of the three *in vitro* candidates (A6, A9, A10; red, purple, and navy respectively) along with the distribution for the newly generated sequences (black). The third plot (Fig 3c) overlays all distributions.

**Figure 3.**
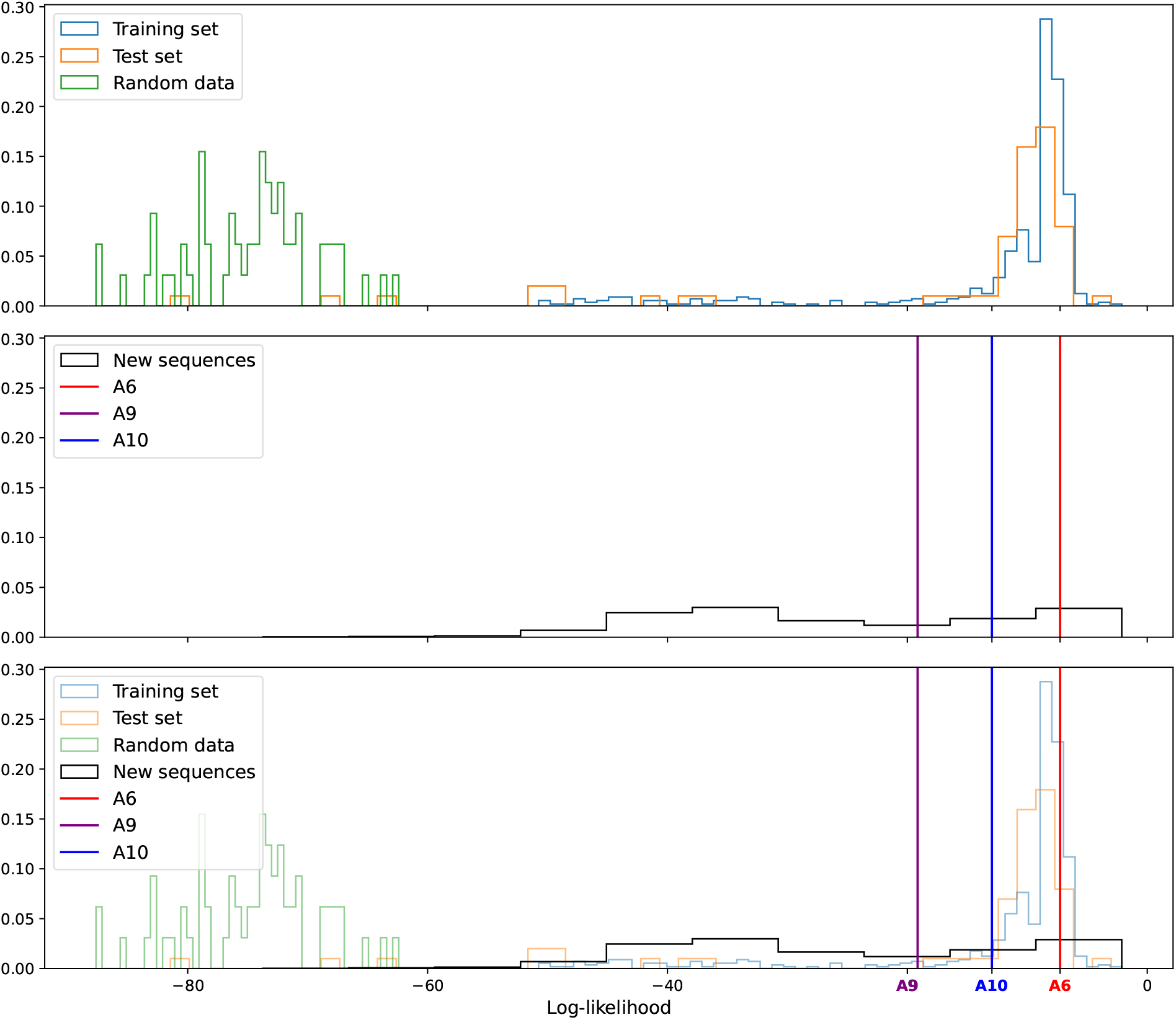
Histogram of log-likelihoods of (a) the observed data set, (b) the generated data set highlighting the in vitro candidates, and (c) their superposition.

**Figure 4.**
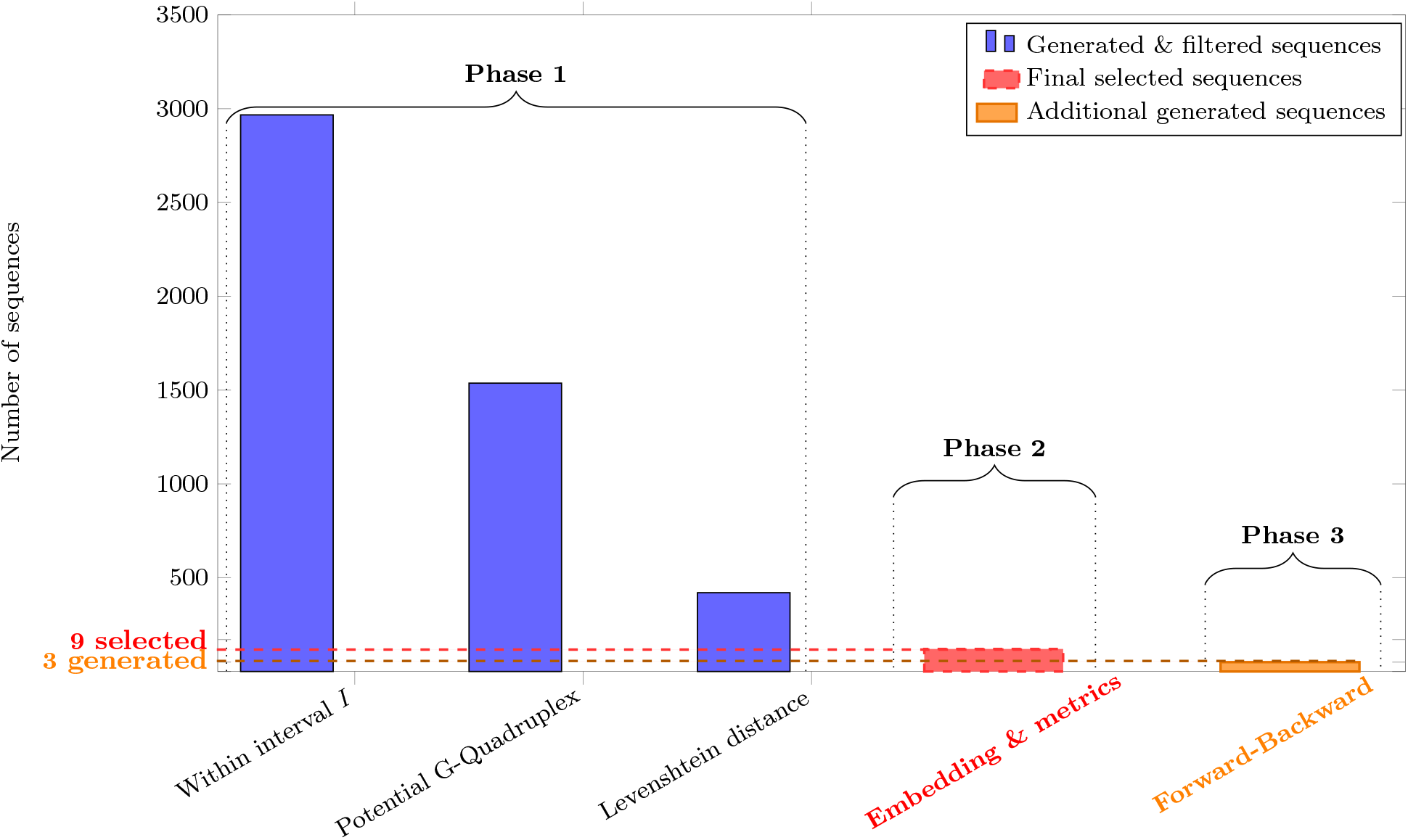
Selection of in silico candidates through successive filters and a complementary generation process Summary of the selection of *in silico* candidates generated by the RBM, including three additional sequences obtained through an alternative generation method. **Within interval:** Sequences were selected if their likelihood values fell within the range of *in vitro* candidates likelihoods (between the lowest and highest values). **Potential G-Quadruplex:** Only sequences with the potential to form a G-Quadruplex structure were retained. **Levenshtein distance:** Sequences close to binders based on Levenshtein distance were kept. **Embedding & metrics:** The remaining sequences were represented in both continuous and sparse embeddings, and those closest to the *in vitro* candidates were selected based on *ℓ*_2_ distance and cosine similarity. **Forward-Backward:** An additional set of 3 *in silico* candidates was obtained through a complementary generation process, independent of the filtering pipeline.

The newly generated sequences are broadly distributed over the upper portion of the log-likelihood axis, excluding only the lowest values, thus covering a wide range beyond the mode of the training and test distributions.

### 3.2. In silico candidates

Our selection procedure is organized into three main phases. Phase 1 applies three successive filtering steps based on log-likelihood, G4 motif detection, and Levenshtein distance. Phase 2 involves choosing a specific combination of sequence representation and distance metric to perform the final selection of nine candidate sequences. Phase 3 complements this approach by generating three additional in silico candidates through an alternative method that directly proposes new sequences, without applying the multi-step filtering used in the previous phases.

#### Phase 1: Filtering with Log-Likelihoods and Levenshtein Distance

Initially, 10,000 sequences are generated according to the retained RBM model.

1. **Initial Generation and Filtering by Log-Likelihood:** To focus on sequences with potential, we filter them by retaining only those whose log-likelihood values fall within the interval 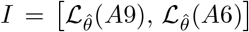, defined by the *in vitro* candidates with the lowest and highest log-likelihoods (see Equation (4)). It is important to note that sequences with high probabilities (or log-likelihoods) correspond to those that appear frequently^6^ in the dataset, but are not necessarily promising candidates. This step reduces the number of sequences from 10,000 to 2,967.
2. **Filtering for Potential G-Quadruplexes:** Next, sequences are further filtered to retain only those likely to contain one or more G-quadruplex motifs. For flexibility, we do not verify G-quadruplex presence using a folding tool but instead we count motifs such as ‘GG’, ‘GGG’, or ‘GGGG’. Sequences with at least four occurrences of any of these motifs are retained, reducing the candidates from 2,967 to 1,537. This filtering ensures a high probability of containing G-quadruplexes without strict confirmation.
3. **Levenshtein Proximity Filtering:** The remaining 1,537 sequences are ranked according to their proximity to each *in vitro* candidate using the Levenshtein distance. From this ranking, we construct a shortlist of 467 sequences from which the *in silico* candidates are eventually selected: 360 one-mutation neighbors (119 per *in vitro* candidate *B* ∈ {*A*6, *A*9, *A*10} plus the *in vitro* candidate itself), 60 obtained as the union of the 20 nearest RBM-generated sequences to each candidate, and 47 obtained as the intersection of the sets of the 350 nearest sequences to A6, A9, and A10. Finally, we ensure that no duplicates are retained in this final pool of 467 sequences.

#### Phase 2: Final selection - Combining dense and sparse representations

In the above filtering steps, only the model (via log-likelihood) and Levenshtein distance were employed. To enhance robustness and validate results, we incorporate *k*-mer sparse representation in conjunction with cosine similarity. We compare four methods based on the combination of intersection or union with sparse or dense representations. Table 3.2 summarizes this comparison, with diagonal elements (highlighted) indicating the combinations selected based on our criteria.

**Table 2.** Comparison of selection methods using sparse and dense representations. Diagonal elements represent the selected combinations.

| | Sparse + Cosine Similarity | Dense + $\ell_2$ Distance |
| --- | --- | --- |
| Union | Selected | Not Selected |
| Intersection | Not Selected | Selected |

Cosine similarity is preferable to Euclidean distance (*ℓ*_2_ norm) for comparing sparse vectors, as it emphasizes the directional similarity of the vectors rather than their magnitude. Unlike Euclidean distance, which overly penalizes differences in magnitude and can distort similarity due to sparsity and dimensionality, cosine similarity inherently normalizes vectors, providing robust and meaningful comparisons in high-dimensional spaces by focusing on shared non-zero dimensions [31, 32]. To move away from the model-based approach that generates and filters sequences based on log-likelihood, we choose the union operation with sparse vectors (k-mers), as this provides more sequences that are closer to the *in vitro* candidate sequences (in terms of cosine similarity) compared to the intersection. Thus, we reserve the intersection filter exclusively for dense vectors.

Out of the 467 pre-filtered sequences, the 9 *in silico* candidates of Phase 2 were selected in two complementary ways:

i. **Union of Sparse Representations (6 sequences):** We first apply cosine-similarity thresholds to retain only sequences sufficiently close to each *in vitro* candidate sequence: cos(*A*6, seq) *>* 0.855, cos(*A*9, seq) *>* 0.92, and cos(*A*10, seq) *>* 0.915. The retained sequences are then ranked by Levenshtein distance to the corresponding *in vitro* candidate sequence, and the two closest to each are kept, yielding a total of six sequences.
ii. **Intersection of Dense Representations (3 sequences):** We then focus on sequences simultaneously close to all three *in vitro* candidate sequences. Only those with a Levenshtein distance ≥ 10 are considered. They are ranked by *ℓ*_2_ distance with respect to A6, A9, and A10, and the intersection of these rankings provides three sequences, ensuring balanced proximity to each *in vitro* candidate sequence.

- Sequences already included in the union are excluded before computing the intersection, in order to prevent duplicates.

#### Phase 3: Direct sequence generation via the RBM Model & negative controls

- **Forward-Backward sampling using the model (3 sequences):** For each *in vitro* candidate sequence *B* ∈ {*A*6, *A*9, *A*10}, we generate *k* vectors ***h***_*i*_ using 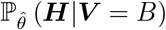, compute the empirical mean 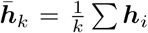, and sample a new sequence ***V*** _new_ using 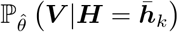. This process is reiterated as needed to ensure a Levenshtein distance of at least 8 from the initial sequence (when *k* is large enough, we often return to the same initial sequence ***V*** = *B*). This choice encourages diversity.
- **Negative Controls (3 sequences):** Negative controls are sequences unlikely to bind targets, generated by reversing the ranking order or using forward-backward models on unfavorable sequences.

### 3.3. Final Outcomes

#### Objective and Method

We propose 12 *in silico* candidate sequences and aim to obtain a visual representation of the different sets of sequences involved in this study: the SELEX-selected sequences, the *in vitro* candidates, the generated and filtered sequences (after phase 1), and the *in silico* candidates. The objective is to assess whether the *in silico* candidates are located close to the *in vitro* ones in the representation space. For this, Principal Component Analysis (PCA) was applied to two types of representations: a dense representation, derived from the RBM model, and a sparse representation, which does not depend on the model.

#### Description of the Figures

Figures 5 and 6 show the PCA projections obtained from the dense and sparse representations, respectively. The same color code is applied to both figures:

**Figure 5.**
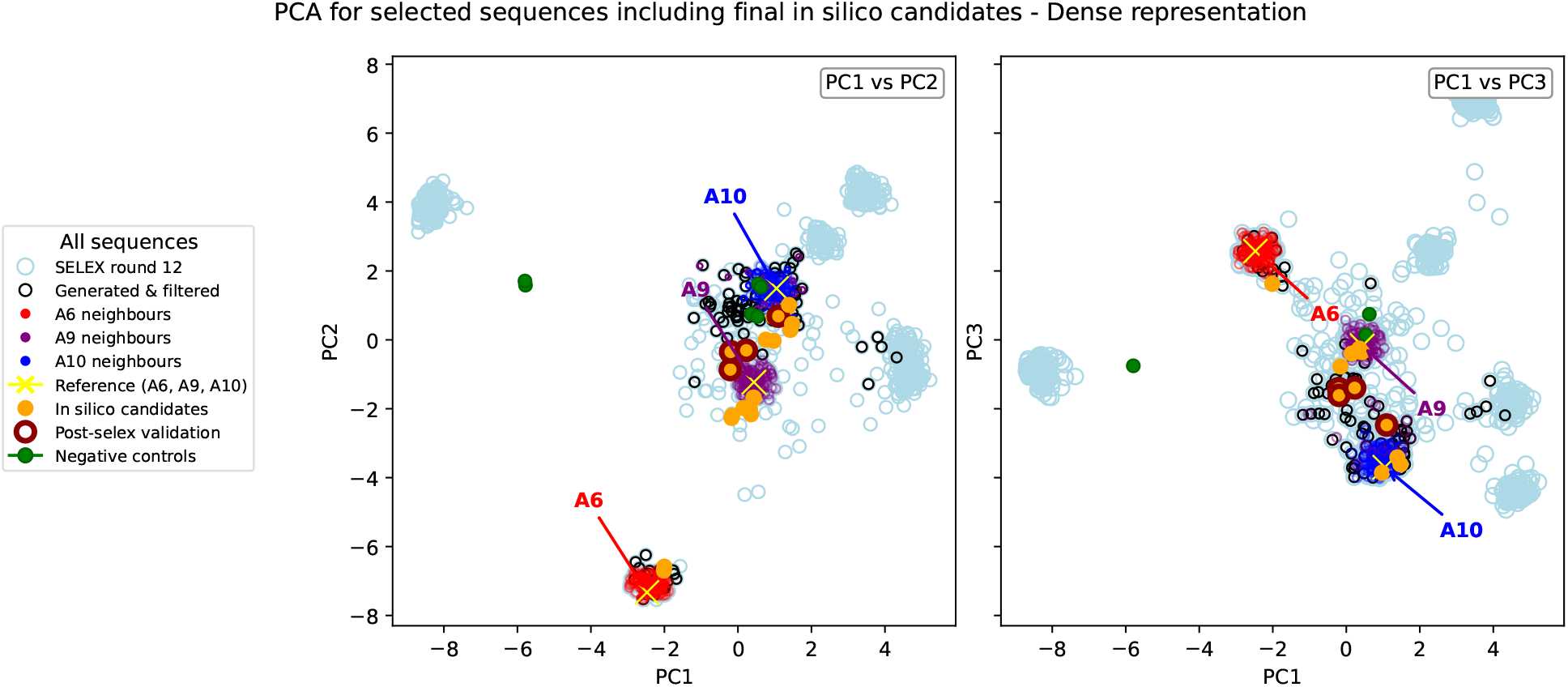
PCA applied on the dense vector representation

**Figure 6.**
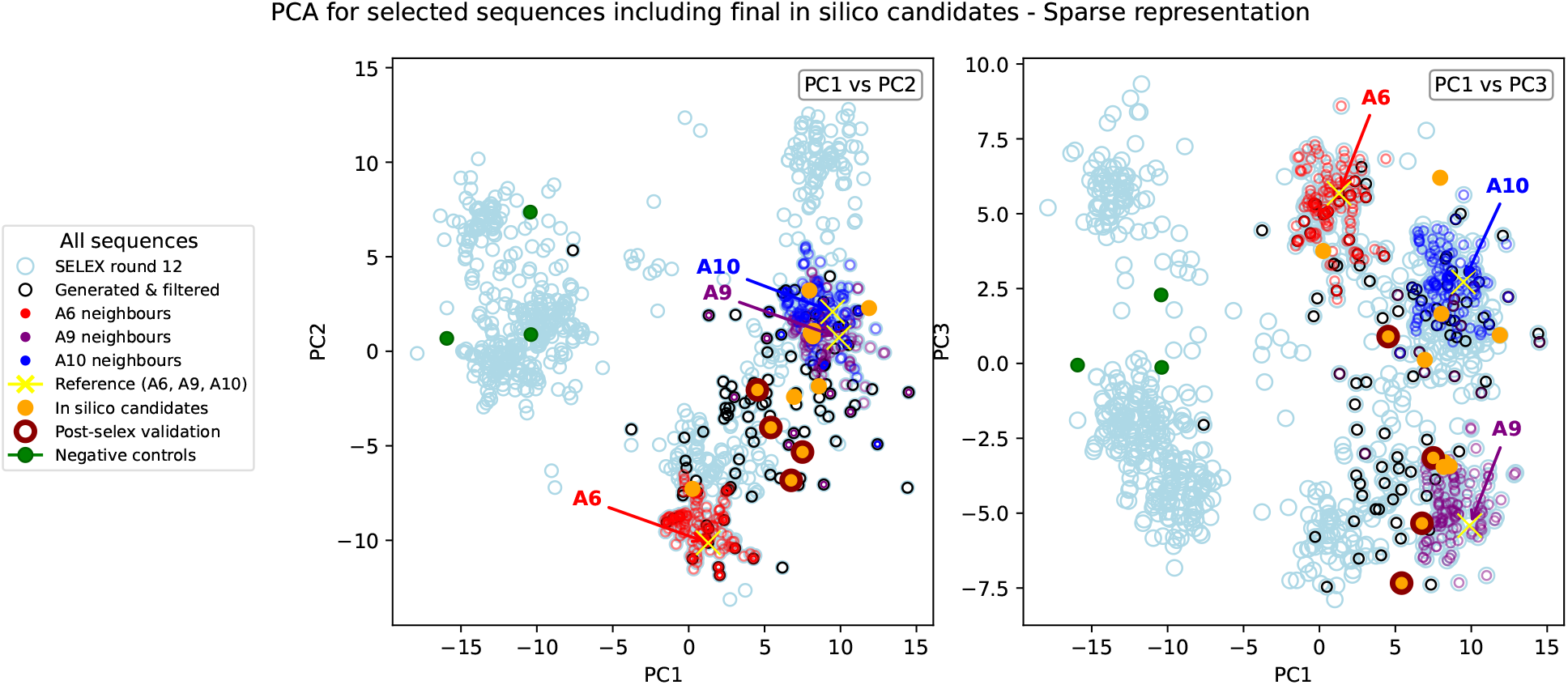
PCA applied on the sparse vector representation

- Light blue dots represent SELEX data from round 12.
- Red, purple, and navy blue dots correspond to sequences A6, A9, and A10 and their neighbors (defined by Levenshtein distance).
- The *in vitro* candidate sequence itself (A6, A9, or A10) is marked with a golden cross.
- Yellow dots represent the 12 *in silico* selected sequences.
- Green dots indicate the 3 negative control sequences.

In Figure 5 (dense representation), the PCA projection reveals three peripheral clusters and one central cluster. The peripheral clusters are associated with the most frequent sequences and their neighbors, while the central cluster includes the remaining sequences, among them A6, A9, and A10. Most of the *in silico* candidates (yellow) lie in the central cluster near the *in vitro* candidates. Two negative controls are also found in this area, whereas the third one is closer to the A1 cluster, which is consistent with its origin from the non-binding sequence A1 (phase 3).

In Figure 6 (sparse representation), a similar overall structure is observed, but A6, A9, and A10 appear closer to each other, particularly A9 and A10. The *in silico* candidates remain located near these *in vitro* sequences, while the negative controls are clearly separated from them.

#### Interpretation and Analysis

For the dense representation, the first three principal components explain about 61% of the total variance, indicating that most of the information is captured by the first few axes and that the representation is compact and informative. Extending to 15 components increases the explained variance to 95%, confirming the good quality of the dense encoding.

For the sparse representation, the results depend strongly on rescaling. Without rescaling, about 79% of the variance is explained by the first six components. After rescaling, the explained variance drops to around 6%, as null components become non-zero and introduce noise, reducing the discriminative power of the PCA.

In terms of spatial organization, both representations show consistent patterns between *in silico* and *in vitro* candidates, confirming their proximity in sequence space. In dense representation, the *in vitro* candidates are slightly more separated, whereas in the sparse representation, they overlap more closely, particularly A9 and A10. Conversely, the negative control sequences appear further from the *in silico* candidates in the sparse representation than in the dense one, suggesting better discrimination in this case.

#### Experimental Validation

Experimental results support the observations made from the PCA analysis. All negative control sequences behave as expected and show no binding signal. Among the 12 *in silico* candidates, four display binding to the target protein with intensities comparable to, or even higher than, those of A6 and A10, while no signal is detected for an unrelated protein, confirming specificity to the target *CspZ* exclusively. Among these four promising *in silico* candidates, three are obtained through the dense vector intersection, and one through the sparse vector union, suggesting that the intersection-based approach may be more effective.

Although the current comparison is based only on *k* -mers of size *k* =1, 2, 3, 4, and 8, the results are encouraging and confirm the relevance of both dense and sparse representations. Further measurements of the dissociation constant (*Kd*) and a detailed analysis of sequence motifs, particularly G-quadruplex structures, are already underway to refine the characterization of these promising candidates.

## 4. Discussion

In this work, we generate new sequences from an RBM trained on the final SELEX round and select promising candidates by comparing them with additional post-SELEX experiments. The overall workflow follows an *in vitro* → *in silico* → *in vitro* loop: SELEX and post-SELEX experiments provide our dataset; the RBM, together with complementary embeddings and distance metrics, is used to model, generate, and rank sequences; and the selected candidates are then tested experimentally under identical post-SELEX conditions.

We choose to analyze sequences solely through their primary structure, which simplifies processing and avoids biases introduced by folding tools. While this approach limits access to secondary and tertiary structural information, it provides a consistent, reproducible representation of the data. Moreover, the hidden layer of the RBM is expected to take into account indirectly this structural information. To complement the dense representation derived from the RBM hidden layer, we introduce an independent, sparse encoding based on *k*-mers. This additional representation offers a way to refine and balance the model-derived features, while maintaining full independence from structure-prediction tools.

Preliminary *in vitro* validation indicates that some *in silico*-selected candidates exhibit a positive response in the first *in vitro* experimental assays.

When compared with the results obtained by Di Gioacchino et al. [20], the model performs well when the training data are sufficiently comprehensive and well balanced. However, when coverage is limited or underrepresented, incorporating *k*-mer-based sparse representations helps reduce model bias and recover relevant information. Treating the hidden layer as a dense embedding further provides a metric space in which distances between sequences can be computed, enabling quantitative comparisons and graphical visualization.

Among the strategies tested for selecting sequences for experimental validation, the most favorable results are obtained using the intersection approach, in which candidates are jointly selected based on their Euclidean proximity in the dense embedding space to the three *in vitro* reference sequences. In contrast, the union-based selection in the sparse space, built from the closest neighbors identified through cosine similarity, as well as the forward-backward generation method, prove less effective. These results indicate that combining complementary selection criteria is more robust than relying on a single representation, although the optimal strategy may vary depending on the dataset.

Building on the present results, an important perspective is to extend this framework by integrating additional sources of information beyond the primary sequences, in particular secondary-structure descriptors when reliable annotations are available. Such an extension could strengthen the sequence representations derived from the RBM and *k*-mer encodings, and provide a more quantitative understanding of the relationships between sequence statistics and binding properties. In addition, applying the proposed pipeline to larger SELEX datasets, as sequencing depth and experimental coverage increase, would enable further improvement in model training and candidate selection, and would help assess the generalizability of the approach across different targets.

## Funding

This work was supported by Sorbonne University program and performed with the support of the Institut des Sciences du Calcul et des Données (ISCD) of Sorbonne University (IDEX SUPER 11-IDEX-0004), of *Centre National de la Recherche Scientifique, of Ministère de l’Enseignement Supérieur et de la Recherche*.

## Supporting Information - Appendix

### 1. In vitro candidates selection

Figure S1 is derived from experimental data, that illustrates the results obtained post-SELEX. For the Dot Blot results, a higher number of “pluses” indicates a higher likelihood of the sequence being favorable, as observed for sequences B3, B4, A6, B7, B8, A9, A10 and A11 (Only in this Figure, were different labels used, consisting of 2–4-character alphanumeric identifiers. abB3, B4, A6, bB7, GST8, A9, A10, and A11).

In the second row, a smaller KD value signifies a more favorable sequence, which is the case for sequences such as B3, B4, A6, and A10. Other criteria include the presence or absence of G-Quadruplexes, among others.

**Table 3.** Characterization of experimental aptamers.

| Aptamères | abB1 | abB2 | aB3 | B4 | GST5 | B6 | bB7 | GST8 | B9 | B10 | A11 | A12 | A13 | A14 | B15 | A16 | CTRL |
| --- | --- | --- | --- | --- | --- | --- | --- | --- | --- | --- | --- | --- | --- | --- | --- | --- | --- |
| Dot BLOT |  |  |  |  |  |  |  |  |  |  |  |  |  |  |  |  |  |
|  | - | - | +++ | ++ | + | +++ | ++++ | ++ | ++ | +++ | ++++ | + | - | - |  |  | - |
| BLI – Bb2<br>(K<sub>D</sub>, nM)<br>(100 mM NaCl) | 565 | 535 | 74,7 | 51,1 | 985 | 49,6 | 7,8 | 702 | 113 | 84,9 | 14,2 | 168 | 975 | 940 | EN COURS<br>D'ANALYSE |  | 1180 |
| Cytométrie<br>de flux | - | - | + | + | - | + | - | - | ++ | +++ | - | + | - | - |  |  | - |

### 2. RBM Optimisation : Contrastive Divergence and Algorithm

#### Contrastive Divergence

Contrastive divergence (CD-k) is commonly used to update the RBM parameters by running a Gibbs chain for a few steps to approximate the log-likelihood gradient (see [33]). However, due to the biased approximation in CD-*k*, alternative methods have been proposed to improve the algorithm. One such approach is parallel tempering [34, 35], an advanced Monte Carlo method that helps reduce bias and improve efficiency in learning RBMs (see [36]). Other techniques include Persistent Contrastive Divergence (PCD) [37], which maintains a persistent Markov chain to generate more stable updates, and Fast Persistent Contrastive Divergence (FPCD) [38], which introduces adaptive learning rates to enhance convergence speed. In Contrastive Divergence, the double expectation over ***v*** and ***h*** is replaced with a point estimate by sampling a single value, 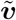. This sample is obtained through Gibbs sampling, initialized at the current training observation, and run for a few steps (*k*) to approximate the true distribution:

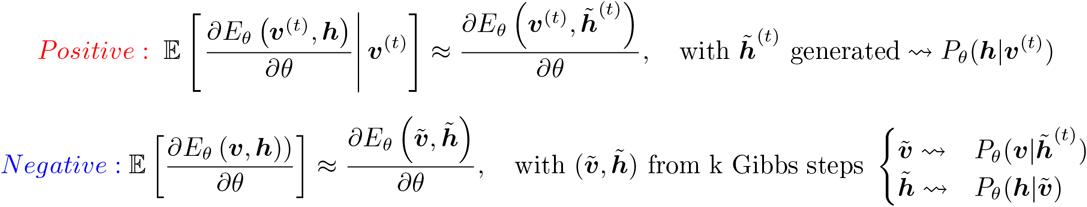

#### Algorithm

The standard practice is to compute the gradient for each sample, average these gradients within each mini-batch, and then update the parameters using the averaged gradient (which, over many batches, is equivalent to averaging over all samples). This approach is adopted in [33, 39, 40, 37]. Although alternative strategies could be explored, such as adding gradients within each batch before averaging across batches, the aim here is not to optimize the internal mechanics of RBMs. Instead, we focus on their potential, such as exploiting the hidden layer for generating neighbors or other downstream applications. Hence, we consider the widely used gradient computation approach to maintain alignment with previous studies [33, 39, 40, 37].

##### Algorithm 1: Training a Restricted Boltzmann Machine

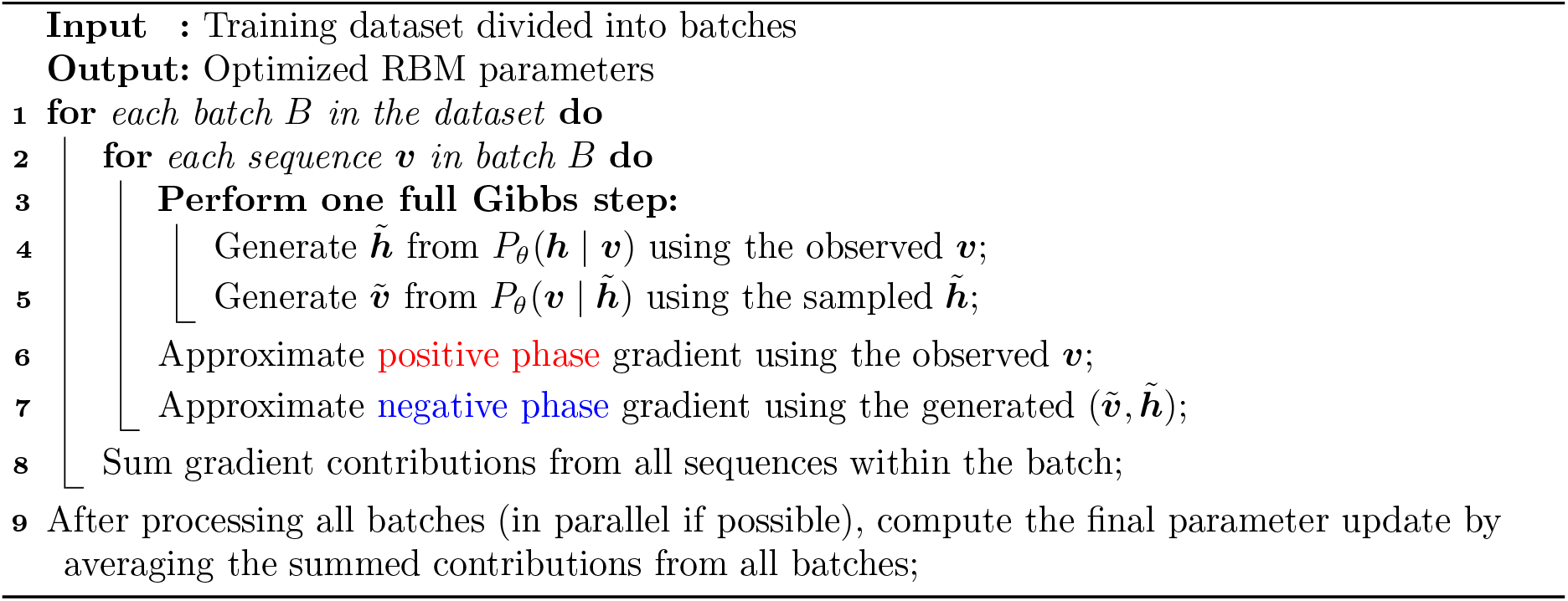

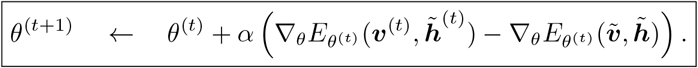

For each observation, we start with ***v***^(*t*)^, we generate 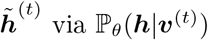 via ℙ *θ* (***h*** | ***v***^(*t*)^) for the positive phase, then perform *k* Gibbs iterations, alternately sampling ***v*** from ***h*** and ***h*** from ***v***, to obtain the final 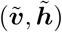 for the negative phase. In general, the bigger *k* is, the less biased the gradient estimate will be. In practice, *k* = 1 works pretty well for pre-training or to extract a good set of characteristics. However, when it comes to evaluation in a test set, it turns out that it is not going that well, the reason is: the model does not investigate for the global minima when using a few steps of Gibbs, while when we do a higher number of steps *k*, we allow the algorithm to investigate farther from the observation.

To simplify the explanation, we have described the algorithm in its standard form, known as the *Vanilla* version. However, in practice, gradient descent for RBMs typically includes two additional terms in Δ*θ* as follows

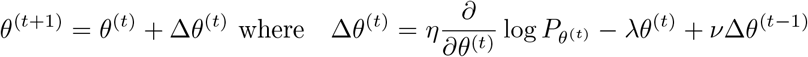

These terms help reduce variance and encourage the algorithm to explore regions that may have high variance but still contribute to effective learning. The term − *λθ*^(*t*)^ comes from the *ℓ*_2_ penalisation of the objective function to get small absolute values for weights. And the term *ν*Δ*θ*^(*t*−1)^ is used to avoid oscillations in the iterative update procedure and to speed up the learning process.

## Footnotes

1 The last row of Table 2.1 represents the entropy for each round, defined as ℍ(*p*) = − ∑_*i*_ *p*_*i*_ log_*N*_ (*p*_*i*_), where *p* = (*p*_1_, …, *p*_*N*_) and *p*_*i*_ denotes the proportion of the i^th^ sequence, *N* is the number of unique sequences in the considered round, and log_*N*_ is the logarithm in base *N*.

2 G-quadruplexes are specific 3D patterns formed by repeated short blocks of the nucleotide G in a DNA sequence. These patterns typically involve quadruplets of consecutive Gs (e.g., GG, GGG or GGGG), which align and fold into stable stacked structures.

3 **batch**: A subset of the training data used to compute a single update of the model parameters during training.

4 **epoch**: One complete pass through the entire training dataset.

5 New uniformly generated sequences

6 By “frequent,” we mean that the group of neighboring sequences (with one or two mutations) is well represented. Even when considering unique sequences, this group appears often because the main sequence has many close variants.

## Notes

### Competing Interest Statement

The authors have declared no competing interest.

### Summary of Updates

We have corrected the author's order and the author's affiliations. Running title has been added. Footnote lines have been removed.

